# An investigation of *Burkholderia* cepacia complex methylomes *via* SMRT sequencing and mutant analysis

**DOI:** 10.1101/2020.03.09.983379

**Authors:** Olga Mannweiler, Marta Pinto-Carbó, Martina Lardi, Kirsty Agnoli, Leo Eberl

## Abstract

The *Burkholderia cepacia* complex (Bcc) is a group of 22 closely related opportunistic pathogens which produce a wide range of bioactive secondary metabolites with great biotechnological potential, for example in biocontrol and bioremediation.

This study aimed to investigate methylation in the Bcc by SMRT sequencing, and to determine the impact of restriction-methylation (RM) systems on genome protection and stability and on phenotypic traits. We constructed and analysed a mutant lacking all RM components in the clinical isolate *B. cenocepacia* H111. We show that a previously identified essential gene of strain H111, *gp51*, encoding a methylase within a prophage region, is required for maintaining the bacteriophage in a lysogenic state. We speculate that epigenetic modification of a phage promoter provides a mechanism for a constant, low level of phage production within the bacterial population. We also found that, in addition to bacteriophage induction, methylation was important in biofilm formation, cell shape, motility, siderophore production and membrane vesicle production. Moreover, we found that DNA methylation had a massive effect on the maintenance of the smallest replicon present in this bacterium, which is essential for its virulence.

*In silico* investigation revealed the presence of two core RM systems, present throughout the Bcc and beyond, suggesting that the acquisition of these RM systems occurred prior to the phylogenetic separation of the Bcc. We used SMRT sequencing of single mutants to experimentally assign the *B. cenocepacia* H111 methylases to their cognate motifs. Analysis of the distribution of methylation patterns suggested roles for m6A methylation in replication, since motifs recognised by the core Type III RM system were more abundant at the replication origins of the three H111 replicons, and in regions encoding functions related to cell motility and iron uptake.

**Author summary:** While nucleotide sequence determines an organism’s proteins, methylation of the nucleotides themselves can confer additional properties. In bacteria, methyltransferases methylate specific motifs to allow discrimination of ‘self’ from ‘non-self’ DNA, e.g. from bacteriophages. Restriction enzymes detect ‘non-self’ methylation patterns and cut foreign DNA. Furthermore, methylation of promoter regions can influence gene expression and hence affect phenotype. In this study, we determined the methylated motifs of four strains from the *Burkholderia cepacia* complex of opportunistic pathogens. Three novel motifs were found, and two that were previously identified in a related species. We deleted the genes encoding the restriction and modification components in a representative strain from among the four sequenced. In this study, methylation is shown to affect various phenotypes, among which maintenance of the lysogenic state of a phage and segregational stability of the smallest megareplicon are most remarkable.

## Introduction

The genus *Burkholderia* is metabolically and ecologically very diverse and consists of bacteria that are able to adapt to and thrive in a wide range of environments, including soil, water, the rhizosphere of plants, in fungi, as well as in association with human and animal hosts (Suarez-Moreno et al., 2012; Coenye et al., 2003). The genus *Burkholderia* was recently divided into two major clades, the *Burkholderia sensu stricto*, containing the Bcc, the *pseudomallei* group and the plant pathogenic species *Burkholderia plantarii, Burkholderia glumae* and *Burkholderia gladioli*, and the newly introduced genera *Paraburkholderia, Caballeronia* and *Robbsia* [1, 2]. These novel genera are usually referred to as *Burkholderia sensu lato* (*Burkholderia* in the broad sense).

The ability of the *Burkholderia* to thrive in highly varied niches is attributed to their unusually large multireplicon genomes, with sizes ranging from 6.2 to 11.5 Mbp [3, 4]. All *Burkholderia sensu stricto* species harbour a primary replicon (chromosome 1, C1) encoding genes with essential housekeeping cellular functions, such as DNA replication, cell division and gene transcription, and a secondary chromosome (chromosome 2, C2), which also carries essential genes. Genes encoded on C2 are less conserved throughout the *Burkholderia*, but are important for niche adaptation [5]. In addition, *Burkholderia* members often carry further highly variable accessory replicons with unique metabolic capabilities. Within the genus *Burkholderia sensu stricto* is a group of closely related bacteria known as the *Burkholderia cepacia* complex (Bcc). The Bcc was originally considered to have three chromosomes, but more recent work has succeeded in curing the third chromosome from all Bcc members in which this was attempted, showing that this replicon is a megaplasmid, rather than a true chromosome. Although this replicon, pC3, does not contain essential genes, its maintenance is important, and is encouraged by various means, such as toxin-antitoxin (TA) systems [6].

Since pC3 is the most variable replicon among Bcc members, and specifies traits such as antifungal activity and in some strains pathogenicity, we thought to use a ‘replicon shuffling’ approach to match the most appropriate pC3 to a ‘chassis’ strain for given applications, to give a plant-beneficial, anti-fungal strain with negligible pathogenicity. Once a Bcc member has been cured of pC3, it is possible to replace it with another pC3 from a different Bcc member into which an origin for conjugal transfer has been inserted. This had been achieved amongst a small number of *B. cenocepacia* strains, and between some *B. cenocepacia* strains and *B. lata* 383. However, all other such transfers attempted between Bcc species were unsuccessful [7]. This stimulated us to investigate potential causes for our difficulties in effecting pC3 transfer, and of these causes, the one we considered most important was defence against incoming foreign DNA by restriction modification (RM) systems.

RM systems utilize DNA methylation as a means of discriminating an organism’s own genome from invading DNA, for example incoming viral or plasmid DNA. Bacteria use methyltransferases to methylate their own genomes, while corresponding restriction endonucleases cleave differently methylated or unmethylated (incoming) DNA [8, 9]. Four different types of RM system have been described based on their subunit composition, cofactor requirements, sequence recognition and cleavage position, known as Types I-IV [10]. In addition to the basic function of RM systems in genome defence, methylation of bacterial genomic DNA is known to have important roles in chromosome replication, DNA mismatch repair and DNA-protein interaction, as well as in the establishment of different cell phenotypes through phase variation. Epigenetic phase variation is a reversible process by which changes in methylation of the DNA result in the silencing or expression of genes [11-14]. Even though both adenine (m6A) and cytosine (m4C and m5C) methylations are present in bacteria, the methylation of adenine bases has been suggested to have greater impact on bacterial gene regulation, whereas cytosine modification has greater impact in higher eukaryotes [11].

DNA methyltransferases have been mostly described as part of RM systems. However, solitary or ‘orphan’ methyltransferases (lacking a cognate restriction enzyme) have been identified in several bacterial species, where they serve various functions. In *E. coli*, deoxyadenosine methylases (Dam) have been found to regulate several important cellular processes like DNA replication, DNA repair and regulation of gene expression [15-17]. Another orphan methyltransferase, CcrM, which was originally identified in *Caulobacter crescentus*, is widely distributed among the *Alphaproteobacteria*. This methylase contributes to the cell cycle control of DNA replication and is essential for *Caulobacter* viability [18, 19]. So far little is known about the impact of DNA methylation on cellular processes in members of the genus *Burkholderia*.

Recent advances in DNA sequencing technology, such as single-molecular, real-time (SMRT) sequencing, have provided new opportunities to detect and analyse the frequency and distribution of methylated bases [20, 21]. Here, we used SMRT technology to detect motifs and modifications in several Bcc members, as well as to investigate methylation patterns in *B. cenocepacia* strain H111. We also deleted the genes encoding putative H111 RM systems to allow us to draw conclusions on the role of RM systems in gene regulation and their impact on bacterial phenotypes.

## Results

### *In silico* comparisons reveal the presence of two core RM systems present throughout the *Burkholderia cepacia* complex

To identify putative RM systems present in our model strain *B. cenocepacia* H111, REBASE, a comprehensive database for DNA restriction and modification curated by NEB was used (http://REBASE.neb.com/REBASE/REBASE.html). Inspection of REBASE showed that seven putative RM system loci are present on chromosomes 1 and 2 (two RM system loci, 3 orphan methylase-encoding loci and 2 restriction endonuclease loci, see Fig 1), while none was identified on megaplasmid pC3 (Fig 1).

**Figure 1.**
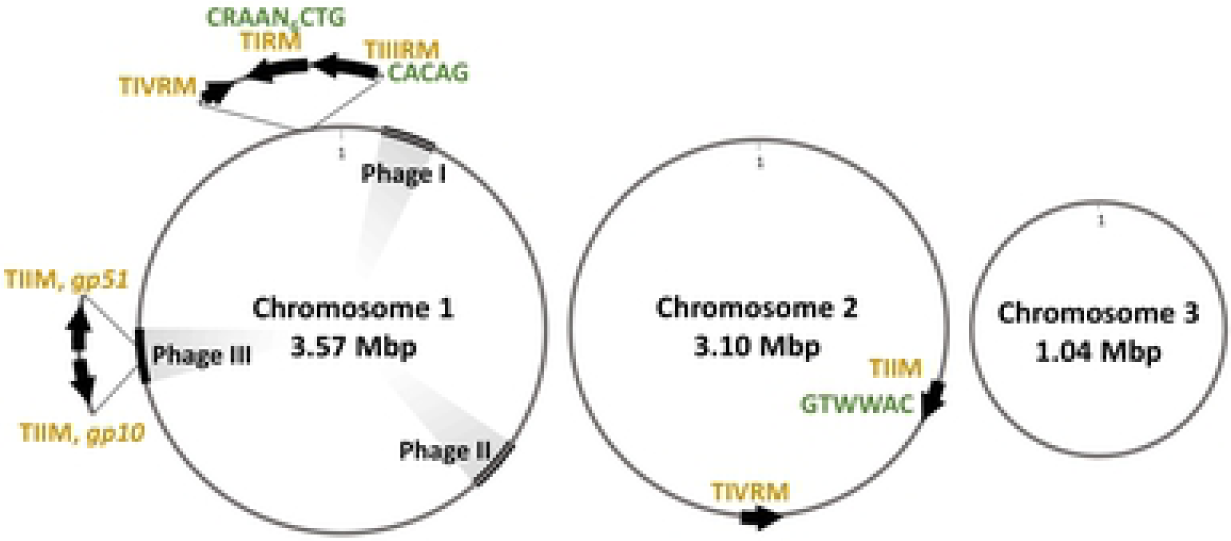
RM systems present in the *B. cenocepacia* H111 genome. RM – restriction and modification system, RE – restriction endonuclease, M – methyltransferase (orphan). Recognition motifs confirmed in this study have been shown in green. Location of prophages has also been shown.

We next compared the RM loci of H111 to those of all other *Burkholderia sensu latu* members entered in REBASE. This comparison revealed that homologues of one H111 RM system (the C1 encoded Type I system) and two orphan methylases (the TII orphan methylase on c2 and *gp10*) were present in numerous strains within the *Burkholderia sensu latu* genomes, while the remaining five RM components present in H111 were very rare. Upon inspection of the strains bearing each RM component, it was apparent that the orphan TII methylase on H111 c2 was present in the majority of *Burkholderia sensu lato* strains, while the H111 TIII RM system was broadly present across the Bcc and the *pseudomallei* group (Fig S1). The prevalence of these RM components in the Bcc led us to consider them as ‘core’ RM components in this complex. These two methylases were previously identified in the *B. pseudomallei* 982 genome, and predictions had been made for their recognition motifs [22].In order to look for any phylogenetic relationship between the *Burkholderia sensu lato* and the RM components homologous to those present in H111, one representative strain was taken from each species present in REBASE, in addition to two commonly studied *B. cenocepacia* strains (J2315 and HI2424), to allow investigation of the diversity of RM components within the species. Since many more strains had been sequenced from the pathogenic *Burkholderia sensu stricto* than the other members, limiting the genomes considered to one per species also reduced bias. As previously mentioned, two additional strains from the species *B. cenocepacia* were included. This allowed a glimpse of within-species differences in RM components. Using representative strains allowed us to consider the entire genome of each representative strain, rather than the amino acid sequences of the RM components identified by REBASE, removing a potential source of error. tBLASTN was used to find homologues of the H111 RM components. These are illustrated as a heat-map in Fig 2, next to a phylogenetic tree of the strains, generated from their concatenated *gyrB* and *rpoD* genes. This analysis highlighted the homology of the TIII RM system (TIIRE and TIIM on Fig2) within the *Burkholderia sensu stricto* (the Bcc, the *pseudomallei* group, and *Burkholderia gladioli, Burkholderia plantarii* and *Burkholderia glumae*). A homologue of the orphan TII methylase (TIIMc2) was found in each of the *Burkholderia sensu lato* representatives, and also within the *Ralstonia pickettii* outgroup, demonstrating the presence of this methylase in the broader *Burkholderiales* order. The phage-encoded *gp10* methylase gene was present within bacteriophage insertions in multiple *Burkholderia sensu lato* species, in a pattern consistent with acquisition by phage transduction. Interestingly, no close homologue of the H111 *gp51* methylase gene, which is part of the same bacteriophage (phage H111-1), was found. The Type IV RE encoded on C1 (TIVRc1) and the TI RM system (TIRM) were specific to *B. cenocepacia* H111, and the remaining TIV RE (TIVRc2) was found in only two other strains.

**Figure 2.**
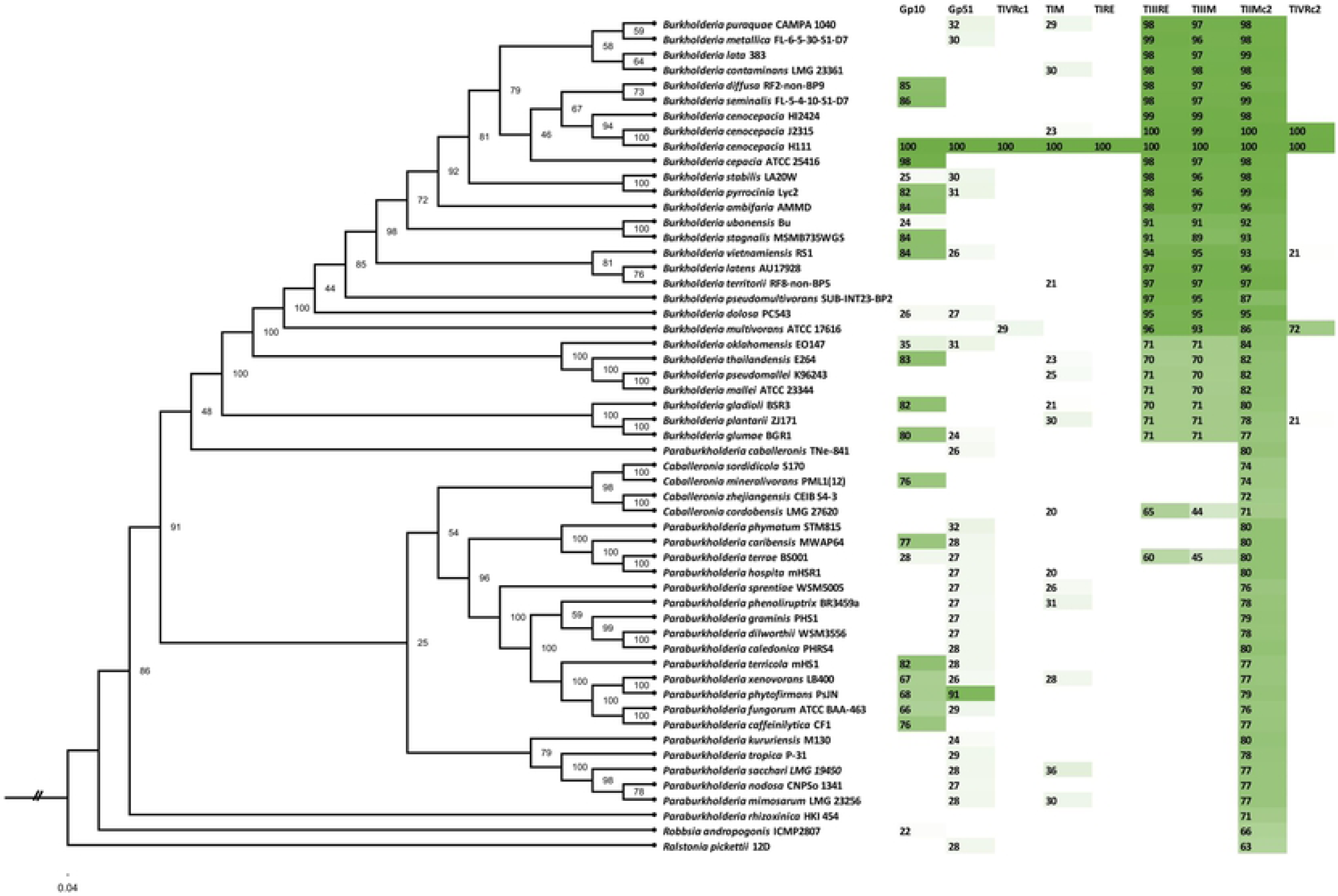
Distribution of homologous of H111 RM components throughout the *Burkholderia sensu lato*. The phylogenetic tree was generated from concatenated *gyrB* and *rpoD* genes, which are essential and highly conserved, with the *Ralstonia pickettii* 12D genes used to give an outgroup. RM components have been abbreviated as follows: *gp51* and *gp10*, type II phage-encoded methylases on C1; TIVRc1, type IV endonuclease encoded on C1; TIRE and TIM make up the type I RM system encoded on C1; TIIIRE and TIIIM, type III RM system encoded on C1; TIIMc2, Type II methylase encoded on c2; TIVRc2, type IV endonuclease encoded on c2. The amino acid sequence of each RM gene was used to query a BLAST database including the genomes of each strain shown. Numbers on the heatmap represent percentage identity, calculated as the number of identical residues between the query and the match, as a percentage of the number of residues present in the H111 query sequence (excluding stop codon). Matches with less than 20 % identity were excluded.

### Verification of predicted sequence recognition motifs and identification of further motifs

To allow us to identify and compare genomic methylation patterns, the methylomes of *B. cenocepacia* H111, *B. lata* 383, *B. ambifaria* AMMD and *B. multivorans* ATCC 17616 were sequenced using Single Molecular, Real-Time (SMRT) sequencing. Various base modifications can be identified through SMRT sequencing due to their specific kinetic signatures, allowing epigenetic studies in different organisms [23].

Our sequencing data confirmed that the two motifs predicted to be methylated in H111 were indeed methylated (CACAG and GTWWAC), and also found methylated motifs for *B. ambifaria* AMMD and *B. multivorans* ATCC 17616, for which no motif predictions had yet been made (Table S1). Furthermore, we identified an asymmetric bi-partite motif in *B. cenocepacia* H111 (5’-CAG-NNNNNN-TTYG-3’) of the type methylated by Type I RM systems [24, 25], of which one was annotated by REBASE, on H111 C1. A Type I motif was also revealed in our *B. multivorans* ATCC 17616 genome sequence. No such Type I RM system was annotated in the publicly available ATCC 17616 genome by REBASE, however inspection of our sequencing data revealed a gene homologous to Type I methylases present in *B. multivorans* D2095 and D2214, which is likely to be responsible for methylating the 5’-CCA-NNNNNN-RTTC-3’ motif.

In addition to the core motifs, we identified another palindromic motif, of the type recognised by II RM systems, in *B. ambifaria* AMMD (RGATCY). [26, 27]. Two such systems are encoded in the AMMD genome. The *B. lata* genome was only methylated at the core motifs, consistent with REBASE predictions which had identified only two methyltransferases in the *B. lata* genome, both close homologues of the H111 core methyltransferases.

### Analysis of H111 methylation patterns suggests importance in cell replication and motility, and iron uptake

In *B. cenocepacia* H111, 60,867 modifications were detected, of which 14,585 were adenine base modifications (m6A) and 2,804 were cytosines (m4C). A total of 14,347 of the detected 14,585 (98%) adenine modifications were assigned to a specific motif, whereas no specific motifs could be identified for the modified m4C bases detected (Fig 3, Fig S2). In addition to m6A and m4C modifications, 43,478 ‘modified bases’ were detected, that showed signatures consistent with some form of modification, but to which the PacBio SMRT Portal could not assign a precise type of modification (Fig 3, Fig S2). These unspecific ‘modified bases’ could represent methylated bases, or alternatively they could result from DNA damage due to stress prior to or during DNA extraction and purification. Such damage also results in modified bases, such as 8-oxoguanine and 8-oxoadenine.

**Figure 3.**
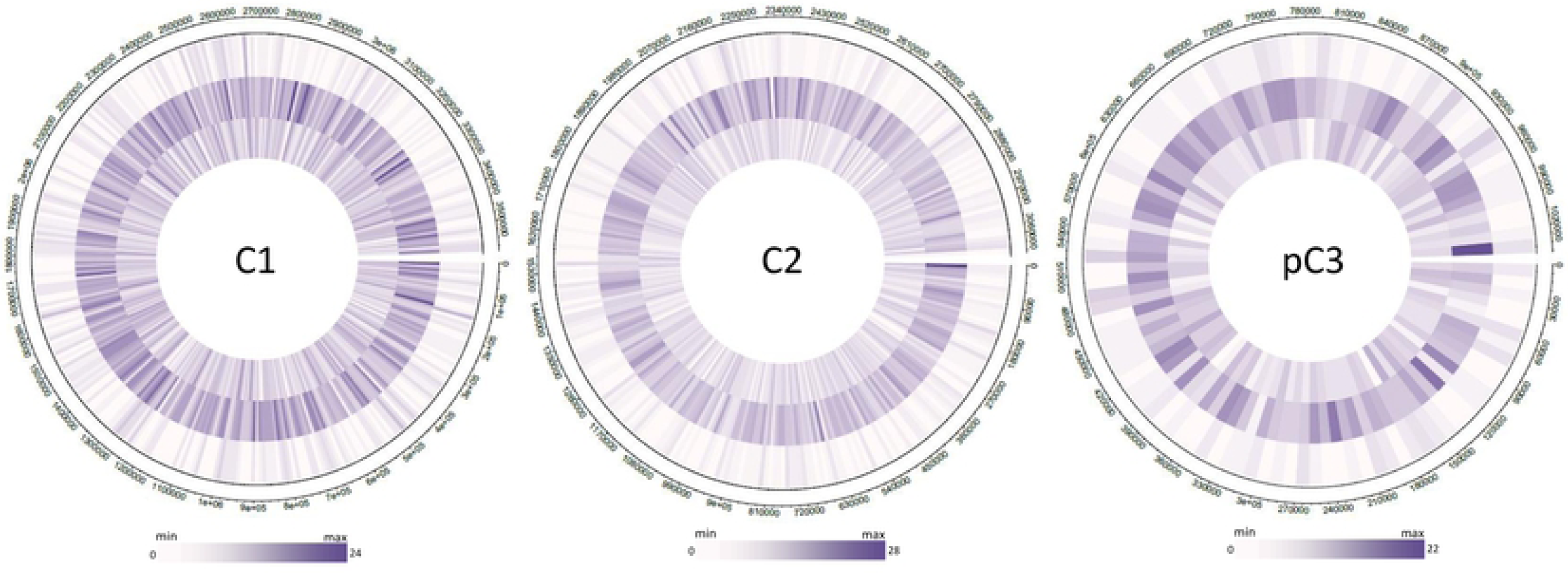
Distribution of m6A base modifications. Patterns in purple represent m6A modifications that were assigned to specific motifs: inner circle motif, CAGNNNNNNTTYG/CRAANNNNNNCTG Type I RM system; second circle motif, CACAG (Type III RM system); outer circle motif, GTWWAC (Type II orphan methylase). Every hatchline in each circle is a representation of a 10,000 basepair window. The numbers of modified bases or motifs per window is represented by the color range. The darkness of colour corresponds to the number of modifications (maximum number of modifications found in a window: C1, 24 modifications; C2, 28 modifications; pC3, 22 modifications). Circles were constructed using the circlize package for R.

To allow us to analyse the H111 methylome as a whole for patterns and hotspots, we divided each replicon’s sequence into 10,000 bp windows, and the abundance of modifications was calculated (Fig 3). We found that m6A and m4C modifications (Fig 3, Fig S2), as well as the uncharacterized ‘modified bases’ were mostly evenly distributed. However, some windows contained an increased number of motifs. Most noticeable was an increase in m6A methylations at the origin of replication of each replicon (Fig 3). These modifications were mainly at CACAG motifs, the motif recognised by one of the core RM systems (Type III RM) identified within the *Burkholderia.* To further analyse the CACAG core RM motif, we evaluated the windows in which this motif occurred most frequently (Table S2). In these hypermethylated windows we observed an abundance of genes that code for proteins involved in cell replication, e.g. cell division protein FtsK (I35_0834, *ftsK*), DNA-directed RNA polymerase (I35_3258), topoisomerase (I35_2384, *parE*), chromosome partitioning proteins ParA and ParB (I35_4003, I35_4004), replication protein (I35_4005) and chromosome segregation ATPases (I35_4006), to name but some. In other hypermethylated windows we observed genes involved in bacterial cell motility and genes coding for transcriptional regulators, transporters, SAM-dependent methylases and proteins involved in iron uptake and utilisation (Table S2).

Windows in which the motif CAG(N)6TTYG/CRAA(N)6CTG (recognized by the Type I RM system) was overrepresented were also scrutinised. Interestingly, this motif was more abundant in a number of genes associated with DNA replication, such as those encoding DNA ligase (I35_2022, *ligA*), DNA gyrase subunit A (I35_0906, *gyrA*) and chromosome partitioning protein Smc (I35_2024, on C1 and pC3). Furthermore, we found genes coding for DNA repair systems (*recCBD*) and other genes associated with DNA repair, such as those encoding the exonuclease family protein YhaO (I35_7871) and an ATPase (I35_2468). Moreover, we observed genes involved in cell motility, such as those encoding MotA, flagellar motor proteins and secretion systems associated with Flp pilus formation where the occurrence of the CAG(N)6TTYG/CRAA(N)6CTG motifs was increased. In addition, genes encoding transcriptional regulators, permeases, ABC transporters, proteins involved in cell shape and cell wall biosynthesis, as well as other membrane proteins, were found in the windows containing a higher than average frequency of methylated CAG(N)6TTYG/CRAA(N)6CTG motifs (Table S2).

The GTWWAC motif was predicted to be recognized by a Type II orphan methylase (one of the core methylases), located within the highly conserved *trp* cluster on chromosome 2. The abundance of this motif was 3.5 to 4-fold lower compared to that of the CACAG motif (the other core motif). The majority of the GTWWAC motifs within the genome were located in regions containing genes coding for transporters, DNA binding proteins and proteins involved in cell motility.

### Construction of an RM null mutant of *B. cenocepacia* H111

To allow investigation of the impact of *B. cenocepacia* H111 RM systems on phenotype, gene expression and genome protection and maintenance, we performed sequential deletions of the seven RM-encoding loci of H111 (two RM system loci, 3 orphan methylase-encoding loci and 2 restriction endonuclease loci, see Fig 1) using an I-SceI-dependent, markerless gene deletion approach [28]. The sequential nature of this deletion strategy resulted in the construction of a series of intermediate mutants in addition to the final RM null mutant, and two additional single RM mutants were also constructed (Table S3). After four rounds of site-directed mutagenesis, it became apparent that the Type II methylase encoded by the prophage III *gp51* gene was essential, in full agreement with our recent mapping of essential genes required for growth of *B. cenocepacia* H111 [29]. For further analysis, we constructed a conditional mutant (strain CM51), in which *gp51* expression was controlled by a rhamnose-inducible promoter (Table S3). This mutant could only grow in the presence of rhamnose, demonstrating that *gp51* is an essential gene (Fig S3). Serendipitously, however, a spontaneous mutant arose which had lost prophage III from its genome, and with it the *gp10* Type II methylase gene and the essential *gp51* Type II methylase gene. This demonstrated that *gp51* was only essential as part of the prophage II region, probably for the maintenance of the phage in a lysogenic state. The remaining methylase gene (*I35_2582*) was deleted by a fifth round of site-directed mutagenesis, resulting in an RM null mutant of *B. cenocepacia* H111. During phenotypic and sequence analysis of this mutant, it was determined that the ∼ 1 Mb megaplasmid pC3 had been lost. A mobilized pC3 was therefore introduced into the null mutant using previously developed techniques [30], to give strain NullpC3^+^. As a control strain for the phenotypic analysis of NullpC3^+^, an analogous version of strain H111 was constructed, which will be referred to as H111pC3^+^.

A further RM null mutant was constructed later, by repeating the final two rounds of mutagenesis, and ensuring that pC3 had been retained at each stage. This unmarked RM null mutant was named newNull and was not used for the majority of the analyses documented here, since no selection could be applied to ensure pC3 maintenance. It should be noted that newNull was examined using the majority of the phenotypic tests used on NullpC3^+^, and gave similar results.

### Verification of loss of methylation in an RM null mutant, and of the methylase cognate to each recognition motif

We used SMRT sequencing to assign the methylated motifs present in the H111 genome to their cognate RM systems. Three methylated motifs were detected in the H111 genome. The genomes of single mutants in the Type I RM on C1 (I35_3254), the Type II methylase on C2 (I35_2582), and the Type III RM system on C1 (I35_1825) were subjected to SMRT sequencing, allowing the predicted motifs 5’-CAG-NNNNNN-TTYG-3’, GTWWAC and CACAG to be confirmed for these methylases, respectively.

The genome of the RM null mutant was also subjected to SMRT sequencing, to verify loss of all methylated motifs by mapping the data from the RM null mutant against the obtained methylome data of the H111 wildtype. No m6A or m4C modifications were detected in the RM null mutant. However, 179,975 bases (∼4-fold higher than in the wild type, in which 43,478 modified bases could not be assigned) were listed as having unassigned modifications by SMRT portal analysis (Fig S2, panel B). While the detected modifications could result from base methylation, it is likely that this represents an increased occurrence of damage to the DNA. SMRT sequencing also verified the complete and clean loss of phage region III from C1, and the loss of pC3.

### Transcriptional profiling by RNAseq shows effects of RM systems on expression of genes involved in cell motility, iron uptake and genome integrity.

As mentioned previously, examination of the hypermethylated windows within the H111 genome had made apparent an increased methylation in and around genes involved in cell division and shape, in chromosome segregation, in DNA repair and in iron uptake and cell motility. In order to examine whether this increase in methylation led to transcriptional and phenotypic effects, and to determine other such effects influenced by methylation, RNAseq and phenotypic analyses were carried out on the H111 RM null mutant, NullpC3^+^, versus its control strain H111pC3^+^.

The unique reads obtained by RNAseq for each replicate of strains NullpC3^+^ and H111pC3^+^ were compared, and the top 500 genes showing the most significant changes in their expression (p-value ≤ 0.01 and absolute log_2_ (Fold change) ≥ 0.5) were taken for further analysis. Of these, 240 were up- and 260 down-regulated in the null mutant compared to the H111 control (Table S4). As expected, the 61 genes of the lost prophage region III, as well as the deleted RM system genes, were among the 260 genes showing decreased expression in the null mutant.

Several genes involved in cell motility were notable in the top 500 most differentially expressed genes. These were *fliI, flgD, fliL, motB_2, motA_1* and I35_1589, encoding the putative fimbriae usher protein StfC, and were less transcribed in the null mutant compared to H111. Also notable was *murA*, which codes for a cell wall hydrolase which plays a role in cell wall formation and cell separation. Another notable gene that showed decreased expression (−0.69 Log_2_ fold change) in the RM null mutant was *trpB*. This gene is part of the tryptophan cluster and is located upstream of the core Type II methylase gene, which was deleted in the construction of the RM null mutant.

Several genes involved in replication, recombination and repair, especially in the SOS response, such as the repressor-encoding gene *lexA*, as well as *recA* and *I35_2899*, the latter of which encodes the putative RecA/RadA recombinase, and the genes *I35_2143* and *I35_2898* coding for the DNA polymerase IV and a homologue of DNA polymerase-like protein PA0670 from *Pseudomonas aeruginosa* [31], involved in mutagenesis were found to be upregulated in the null mutant compared to H111. Other genes important for DNA replication and recombination (*I35_1669*, coding for a homolog of eukaryotic DNA ligase III and *dnaE_2*, encoding a DNA polymerase III alpha subunit), as well as DNA repair (*I35_7256*, coding for an exonuclease subunit A, part of the UvrABC DNA repair system, which catalyses the recognition and processing of DNA lesions) also showed an increase in transcription in the null mutant compared to H111 [32]. In addition, we observed higher read counts of several genes coding for assembly proteins of the two remaining prophages in the null mutant (Table S4). Phage expression in *B. thailandensis* has been shown to be linked to the SOS response [33]. Further, we observed upregulation of genes involved in lipid transport and metabolism (e.g.: *pcaI, pcaJ, I35_1898*, encoding a putative 3-ketoacyl-CoA thiolase and *I35_2250*, encoding a putative Acyl-CoA dehydrogenase family protein). Moreover, genes known to be involved in secondary metabolite biosynthesis, transport and metabolism, especially in pyochelin biosynthesis and utilization (e.g.: *pchB, pchE, pchR, ftpA, ftpB*) were upregulated in the RM null mutant compared to H111. Genes responsible for inorganic ion transport and metabolism (*katB*, encoding catalase/peroxidase and *I35_1307*, putatively coding for cytochrome C peroxidase) also showed up-regulation in the mutant (Table S4).

### Phenotypic analysis of the RM mutant

To further investigate the importance of RM components for the characteristics suggested by genomic methylation patterns and transcriptomic analysis (cell replication, cell morphology, DNA repair, cell motility, iron uptake and maintenance of lysogeny), and in other possible traits, phenotypic assays were carried out. As a basis for these assays, growth in liquid culture of H111pC3^+^ and NullpC3^+^ was examined spectrophotometrically and found to be comparable (Fig S4).

Introduction of megaplasmid pC3 into the RM system null mutant was 24-fold more efficient than into wild type

Given that the accepted main purpose of RM systems is to limit the intrusion of foreign DNA into the genome [34], we tested the RM null mutant for efficiency of replicon introduction by conjugation. As previously mentioned, the third replicon of the Bcc, pC3, can be moved between certain Bcc members, after the integration of an origin for conjugal transfer into pC3 [7]. Transfer efficiency was determined using *B. cenocepacia* K56-2 as the donor, and either the original RM null mutant (NullΔpC3) or H111Δc3 as recipient. A 24-fold higher transfer efficiency rate was observed into the RM system null mutant compared to the H111 wild type, confirming that RM systems play an important role in protecting the genome against incoming DNA (Fig 4, panel A). In view of our initial hopes of establishing a protocol for replicon shuffling within the *Burkholderia* genus, and given the increase in efficiency of conjugal uptake by NullΔpC3, transfer of pC3 from *B. vietnamiensis* LMG 10929 and *B. ambifaria* AMMD into NullΔpC3 was also attempted. This, however, remained unsuccessful.

**Figure 4.**
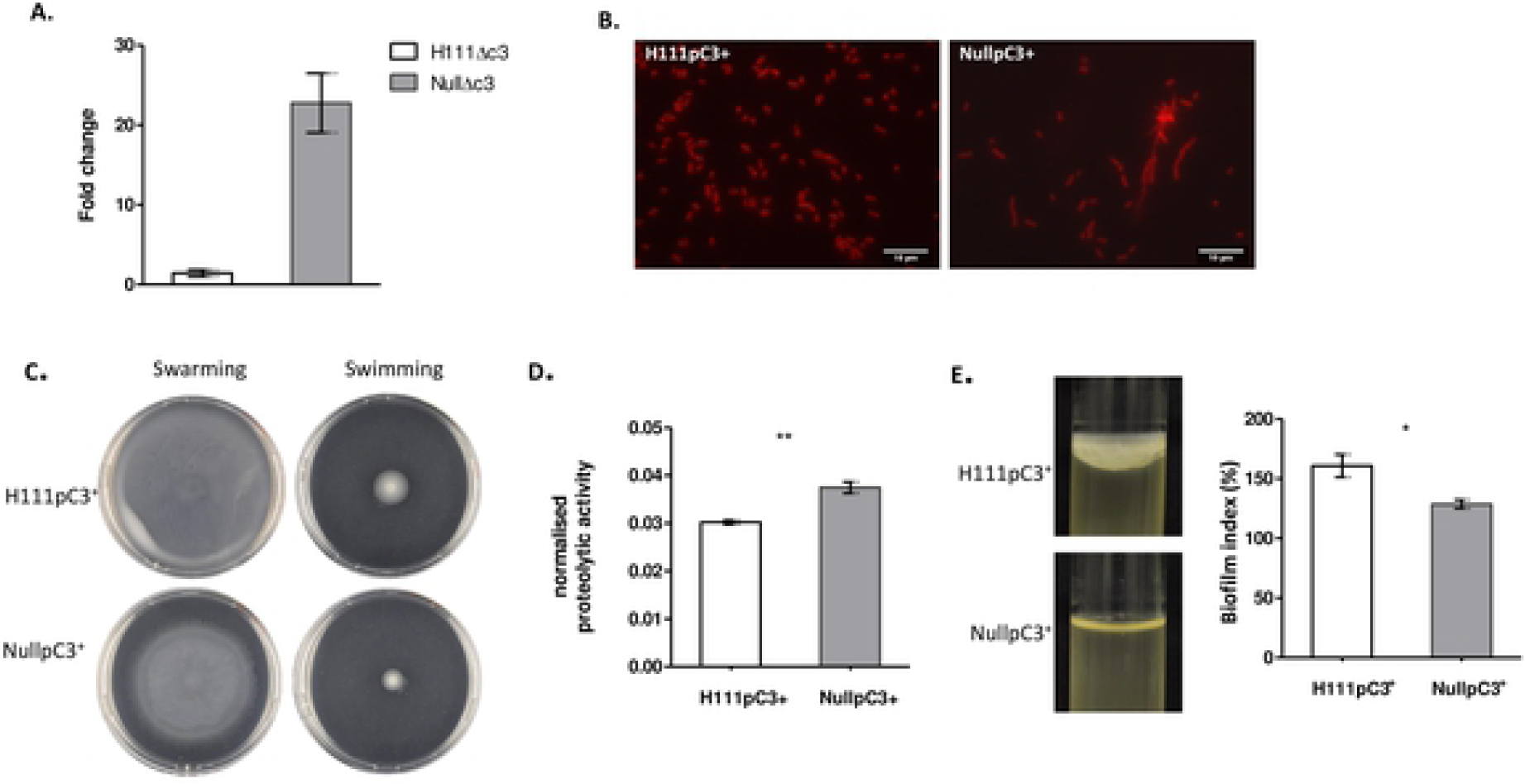
Phenotypic changes observed upon RM system loss. **A.** Conjugal uptake was increased 24-fold in the absence of RM systems. Rifampicin resistant derivatives of pC3-cured strains of NullpC3^-^ and H111△c3 were used as recipients for conjugal transfer. K56-2 pC3 was mobilized by the integration of pSHAFT2, which carries an *oriT* and a chloramphenicol resistance marker. Transfer efficiency was calculated by dividing CFU on media containing Rifampicin + chloramphenicol (total ex-conjugants) by CFU on media containing Rifampicin (total recipient cells). Fold-change in transfer efficiency for H111△c3 and NullpC3^-^ Rifampicin resistant recipients is shown. Error bars represent the standard deviation (SD). **B.** Fluorescence microscopy reveals filamentous cells. Exponential phase cells were subjected to microscopic analysis. Cells were observed with an epifluorescence Leica DM6000 B research microscope at 100 x magnification. **C.** Cell motility is decreased in the absence of RM systems. Representative images from triplicates datasets for swimming and swarming motility are shown. **D**. Proteolytic activity is increased in the RM system null mutant. Bars represent the mean of three technical replicates. Error bars represent the standard deviation (SD). The absorbance at OD_442_ was measured and normalized against the cell density OD_600_. Significance was determined using a two-tailed t-test (p-value: 0.0047). **E.** RM systems are important for biofilm formation. Photographs illustrate differences in pellicle formation, image shown is representative of a dataset of at least 3 replicates. Graph shows biofilm formation using the crystal violet assay. Error bars indicate SD, n=3. Significance was determined using a two-tailed t-test (p-value: 0.0348).

### A decrease in pC3 stability in the RM null mutant confirms the importance of RM systems in genome integrity

To determine the frequency of pC3 loss, a modified version of an experiment previously described in was carried out. The RM system null mutant and H111 control strain were modified to allow positive selection of cells that had lost pC3. To achieve this, the trimethoprim (Tp) resistance gene (*dhfrII*) was placed under the regulation of a modified and tightly controlled lac promotor, and integrated onto C1. The gene coding for the Lac repressor and a gentamycin resistance marker (for selection) were inserted into pC3. As a result, cells bearing pC3 were Tp sensitive, due to repression of *dhfrII* by the Lac repressor. This repression was relieved upon loss of pC3, resulting in colony growth on medium supplemented with Tp. Loss of pC3 in the RM system null mutant strain was 173-fold higher compared to the H111 control strain, demonstrating that RM systems do indeed play a central role in replicon maintenance (Fig 5, panel A).

**Figure 5.**
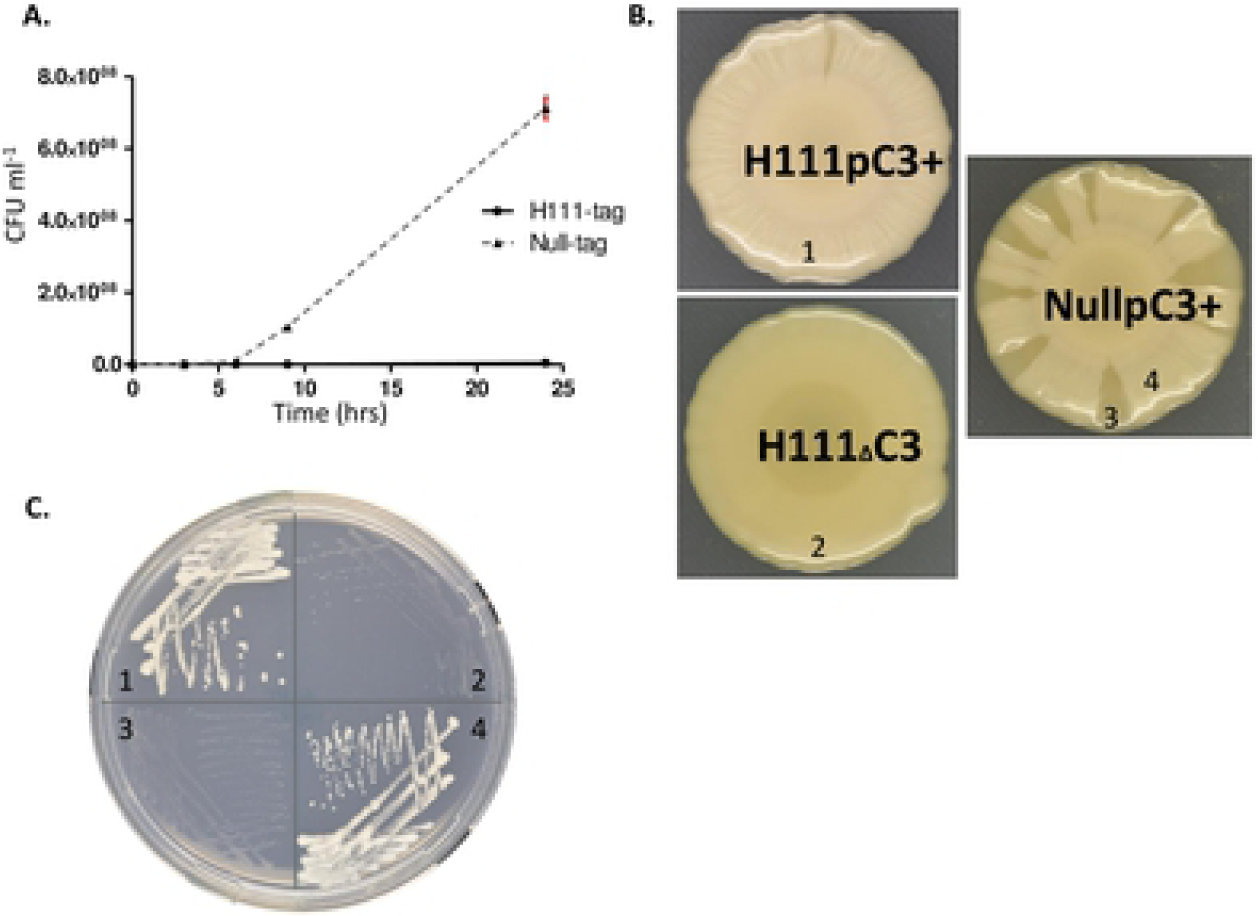
pC3 loss was more frequent in the absence of RM systems. **A.** To assess pC3 stability, strains H111-tag and Null-tag, which were designed to become Tp resistant on loss of the pC3 replicon, were grown in rich medium for 24 hours at 37 °C. Frequency of pC3 deficient cells was plotted. Error bars indicate SD, n=3**. B.** Increased pC3 loss in the absence of RM systems was phenotypically visible on NYG medium. **C.** Inoculum from places indicated by numbers in panel B was streaked on M9 ura medium [5] to confirm the presence/absence of pC3. Strong growth indicates pC3 presence.

This decrease in pC3 stability in the RM null mutant strain was also observed phenotypically. The loss of pC3 from H111 alters the phenotype on NYG plates, as a result of reduced EPS production [5]. Loss of pC3 in the RM null mutant occurred so frequently it could be observed through the formation of wedge-shaped areas with a more transparent appearance (Fig 5, panels B and C). This phenotypic change occurs due to the location of the *shvR* regulator gene on pC3, which is known to influence colony morphology [35]. When sampled and analysed for the presence of pC3 by PCR, the absence of pC3 from the more transparent wedges was confirmed.

### Loss of RM systems leads to filamentous cell growth

Observation of the RM null mutant (NullpC3^+^) and the analogous H111 control strain by fluorescence microscopy revealed that the RM null strain formed cell filaments, while the H111 control did not (Fig 4, panel B). A filamentous phenotype can occur due to the replication arrest caused by the SOS response [36].

### RM systems influence cell motility, biofilm formation and proteolytic activity

As suggested by the observed methylation patterns and our RNAseq data, we observed a reduction in swimming and swarming motility in the RM system null mutant compared to the control strain (Fig 4, panel C). We investigated other phenotypes known to be associated with the RpoN sigma factor, since this was found to be less expressed in the RM null mutant by our RNA-seq analysis. The presence of RpoN is known to repress multiple phenotypes, including the production and secretion of extracellular proteases, EPS, and biofilm formation [37-39]. Interestingly, we saw an increase in protease activity in the null mutant compared to H111 (Fig 4, panel D). There was no difference in EPS production between the two strains (Fig S4), however biofilm formation was significantly decreased in the RM null mutant (Fig 4, Panel E).

### RM systems are not involved in oxidative, heat, membrane damage or osmotic stress tolerance, or in antifungal activity

Tests of the RM null mutant vs the H111 control strain for persistence under oxidative, osmotic, membrane damage and heat stresses showed no significant difference between the two strains (Fig S5). Tests for antifungal activity and pathogenicity against wax moth larvae likewise showed no differences (Fig S5 panel E).

### Deletion of RM systems leads to an increase in phage and membrane vesicle production

Transmission electron microscopy (TEM) was used to investigate the presence of phages and phage-like structures in the supernatants of H111 wild type and the clean RM system null mutant, which our RNAseq analysis suggested was increased in the RM null mutant. In the electron micrographs, we observed the presence of phages and phage tails, either from partially assembled phages or tailocins (bacteriocins) (Fig 6). Furthermore, we discovered long fibres indicating flagella, and other tube-like structures of unknown function. We also observed membrane vesicles of varying sizes. To compare vesicle production between the wild type and the null strain, membrane vesicles were collected and quantified by staining with FM 1-43 fluorescent dye, which binds the cell membrane. We observed a 2.2-fold increase in MV production in the RM null mutant compared to the H111 wild type (Fig 6 panel C), probably as a result of phage-triggered cell lysis (Turnbull et al., 2016).

**Figure 6.**
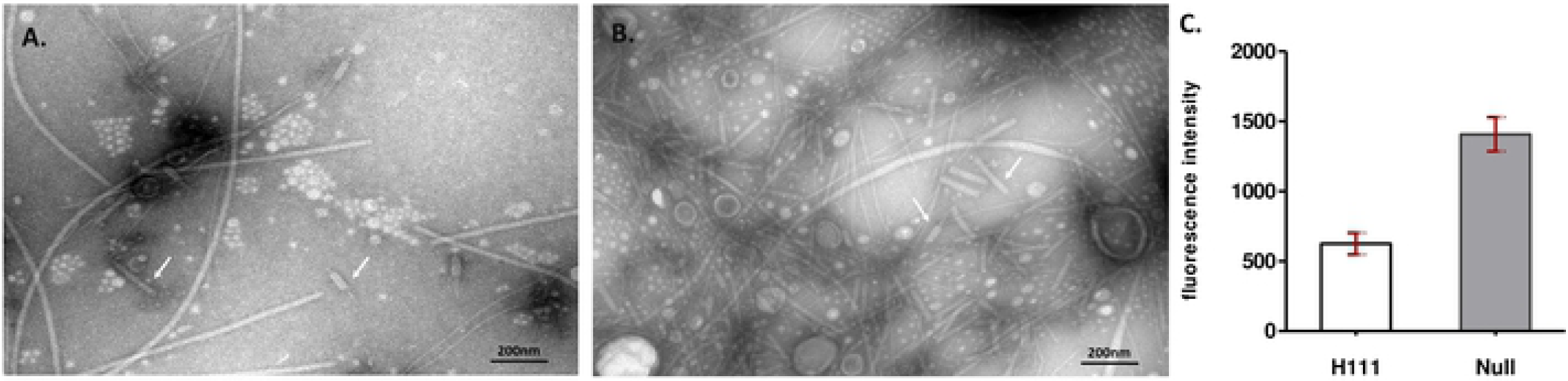
Membrane vesicles (MV) and phage particles were more abundant in cultures of the RM null mutant. Concentrated culture supernatants were visualised at 180,000X magnification by TEM. White arrows indicate phage tails of the *Myoviridae* or *Siphoviridae* family. **A.** H111pC3^+^. **B.** RM null mutant, NullpC3^+^. **C.** Membrane vesicles were more abundant in the absence of RM systems. Concentrated supernatants from the H111 control strain (H111pC3^+^) and the RM null mutant, NullpC3^+^, were stained with FM 1-43, to quantify membrane vesicles (MV). Error bars indicate SD, n=3.

### Chemical and phenotypic assays for pyochelin production support methylation pattern observations and RNAseq results

Genes involved in the biosynthesis and utilization of the siderophore pyochelin showed a transcriptional increase in the RM system null mutant compared to H111 (Table S4). This was investigated phenotypically using CAS plates, in which iron bound to CAS dye can be scavenged by siderophores, resulting in a clearer halo around the colony tested. There was no clear difference in halo size between the RM null mutant and the H111 control strain (result not shown), perhaps because overall siderophore production was similar between the strains as a result of a decrease in expression of the ornibactin genes *orbK* and *orbE* observed in our RNAseq analysis. We grew strains NullpC3^+^ and H111pC3^+^ in an iron limited medium and extracted the siderophores produced. The extracted siderophores were separated by thin layer chromatography (TLC) and visualised by dipping the plate into an FeCl_3_ solution. This gave rise to brown bands, indicating the location of iron chelators on the TLC plate. The bands observed for the RM null mutant sample were clearly darker than for the H111 control strain, suggesting that this strain produced more pyochelin than the control. LC-MS was used to identify the pyochelin fraction, and the relative amount of pyochelin was approximated at around two-fold higher in the RM null strain by determination of the area under the LC-MS curves (Fig 7).

**Figure 7.**
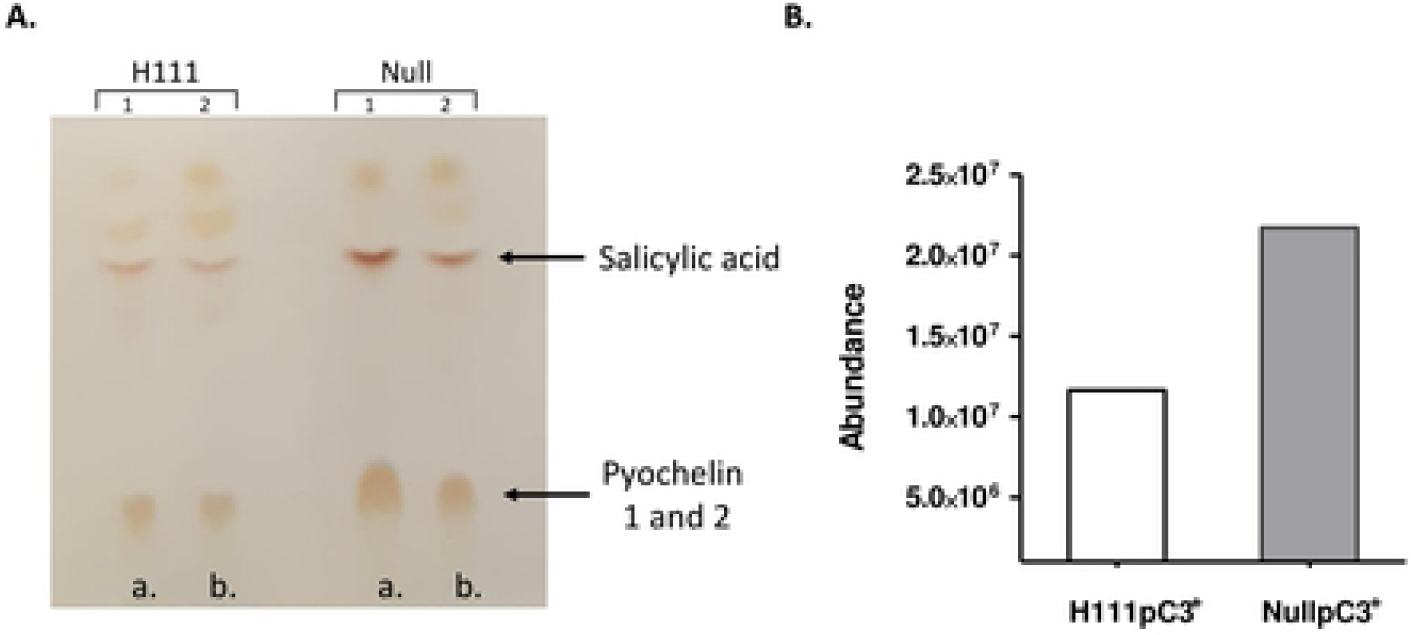
The absence of RM systems results in an increase in pyochelin production. **A.** Thin layer chromatography performed on the RM system null mutant and H111 control suggests that pyochelin production is increased in the null mutant. Siderophores were extracted and separated by TLC. Chloroform-acetic acid-ethanol at 90:5:10 [vol/vol] was used as the developing solvent. Ferric chloride was used to visualize the siderophores. LC-MC/MS confirmed the location of pyochelin and salicylic acid on the TLC plate. LC-MC/MS carried out with Δppm < 3 ppm. **B.** Estimation of pyochelin content. Replicates marked with (b.) in panel A were used. Differences in pyochelin production were approximately quantified by determining the area under the LC-MS curve.

## Discussion

DNA methylation is important for various bacterial cell functions, including host defence, genome integrity and regulation of cellular processes [18, 40]. In this study, we aimed to investigate the methylome of *B. cenocepacia* H111, to allow us to identify specific methylation patterns as well as to study the effects of epigenetics on a broad range of biological processes. We identified a core RM system (*I35_1826, I35_1825*) and a core orphan methylase (*I35_2582*) *via in silico* analysis, and found during *in silico* analysis that the former was conserved throughout the *Burkholderia sensu stricto*, while the latter was present throughout the entire *Burkholderia sensu lato* and beyond, suggesting the acquisition of these components occurred before phylogenetic separation of these clades, and implying their importance since there appears to be strong selective pressure for their maintenance. It should be noted that the core Type II methyltransferase is part of the region of C2 containing the majority of this replicon’s essential genes (BCAM0911 to BCAM0995 in *B. cenocepacia* J2315, I35_2579-I35_2673 in H111). It has been speculated that the movement of this cluster from the primary chromosome, where it is found in the closely related genus *Ralstonia*, to a plasmid might have been a key occurrence in the separation of the genera *Burkholderia* and *Ralstonia* [29].

We also found that three RM systems are present on H111 C1 that are very specific to this strain. Close homologues (defined for our purposes as having a percentage identity of at least 80 %) of the methylase of the Type I RM system were found in six strains in addition to H111, while close homologues of the Type IV endonucleases encoded on C1 and C2 were present in one and six additional strains respectively. This suggests that when we talk of RM systems preventing the successful transfer of ‘foreign’ DNA, even highly related strains are included.

To investigate the distribution of methylated bases, we made use of single-molecular, real-time (SMRT) sequencing technology, to reveal the extent of methylation within the genomes of four Bcc members, each from separate species. Our sequencing data confirmed the two core methylated motifs predicted by REBASE (CACAG and GTWWAC), in the strains sequenced. In addition to the core motifs, further motifs were found in three of the four sequenced strains, while *B. lata* 383 showed only the core methylated motifs, consistent with REBASE predictions. The methylated motifs identified in *B. cenocepacia* H111 were later experimentally assigned to their cognate methylases by SMRT sequencing of single mutants.

We observed that the methylated CACAG motif occurred more frequently in and around the origins of replication of the three H111 replicons. Studies in other *Proteobacteria* have shown that DNA methylation is important for regulation of chromosomal replication, and that m6A modification, for example of the GATC motif in *E. coli*, is densest at the origin of replication (*oriC*) [18, 41]. Furthermore, we evaluated the windows in which CACAG, recognized by the Type III methylase, and the motif CAG(N)6TTYG (recognized by the Type I methylase), were most frequent. The type of genes present in these hypermethylated windows suggested that these methylases might be particularly involved in the regulation of genes involved in DNA replication and repair, and in cell motility. This was corroborated by the increase in SOS gene transcription in the null mutant, as evidenced by RNAseq, and the decrease in motility observed phenotypically in the RM null mutant. The Type II core orphan methylase located in the highly conserved tryptophan operon of H111 recognises the motif GTWWAC. We speculated that the presence of this methylase might be involved in the regulation of the *trp* genes, required for tryptophan production, indeed the RM null mutant showed a decrease in expression of the *trpB* gene in our RNAseq analysis. A single gene deletion mutant of this Type II orphan methylase (SM-OMC2) did not show significant differences in growth in the presence and absence of tryptophan. However, we did observe the increased production of a brown/orange pigment, presumably melanin, when growing the mutant strain in nutrient rich IST medium. Melanin can act to quench reactive oxygen species [42, 43], and therefore its production could reflect an increase in intercellular stress. Interestingly, this methylase was found to be highly conserved throughout the *Burkholderia sensu lato*, and even beyond, suggesting that it was acquired early in evolution. While its high level of conservation might reflect selective pressure for maintenance of the methylase, it might also occur due to the previously mentioned essential nature of the region surrounding it, as the gene cluster from I35_2579 to I35_2673 was previously shown to contain the majority of the essential genes present on C2 [29].

To investigate the impact of the RM systems on the phenotypes of H111, we sequentially deleted the RM systems and components present in its genome. Attempts to delete the orphan methylase encoded by *gp51*, followed by the construction of a conditional mutant, revealed that this methylase is essential. This suggests that the encoded methylase is important in maintaining the lysogenic state of the phage, as has been previously demonstrated for the Dam methyltransferase in enterohemorrhagic *E. coli*, which carries the Shiga toxin-encoding bacteriophage 933W [44]. Gene expression can be altered by promoter methylation, which in most cases prevents expression of a gene. The briefly occuring hemi-methylation of replicons following replication can allow expression of such genes. The essential role played by *gp51* in lysogeny leads us to speculate that by linking induction with the cell cycle via methylation, ϕ1 is able to ensure a constant, low level of induction by an epigenetically triggered mechanism that creates a stochastic switch.

Our null mutant exhibited an increased transcription of genes involved in SOS response, which triggers phage induction [33, 45],. It is interesting to note that in our *in-silico* screen for homologues of the H111 RM components within the *Burkholderia sensu lato*, we found that while homologues of *gp10* were frequently present, *gp51* was very rare. It therefore appears likely that the Gp51 methylase has the ability to confer protection from restriction endonucleases upon entering a new bacterial host [17].

We used transmission electron microscopy (TEM) to investigate the presence of phages and phage-like structures in the supernatants of H111 and the RM null mutant. We detected an increase in phage-like structures in null mutant supernatants, confirming the observation made in the RNAseq analysis. It should be noted that this occurred despite the loss of phage region III, which encodes ϕH111-1, the only confirmed active bacteriophage of H111. In addition, we noticed the presence of membrane vesicles (MV) of varying sizes, and upon analysis found a 2.2-fold increase in MV production in the RM system null mutant compared to the H111 wild type. Toyofuku and colleagues recently showed that an increase in prophage-encoded endolysin triggers MV formation in *P. aeruginosa* and *Bacillus subtilis* [46, 47].

The RM system null mutant was sequenced to verify loss of all methylated motifs. We confirmed the loss of all m6A and m4C modifications previously detected in the wild type. However, compared to the wild type strain, the abundance of unassigned modified bases was 4-fold higher in the null mutant than WT. DNA methylation slows base incorporation in SMRT sequencing, but so too does DNA damage. The increase in unassigned modifications is likely to represent increased nicks in the genome sequence due to DNA damage. Various genes involved in replication, recombination and repair, especially in the SOS response, such as *lexA, recA*, genes coding for DNA polymerase IV (which acts during the SOS response) and an exonuclease subunit A (part of the UvrABC DNA repair system), were found to be upregulated in the null mutant compared to H111, suggesting that the RM null mutant might be subject to a higher level of DNA damage. In *E. coli*, Dam-mutants are subject to increased transcription of the SOS regulon. This effect is thought to occur indirectly; in the absence of Dam methylase-mediated strand discrimination, the mismatch repair system (MutHLS) causes dsDNA breaks, leading to SOS regulon induction [48, 49].

We observed several phenotypic changes in the RM null mutant that are known to be associated with sigma factor RpoN, whose encoding gene showed reduced expression in out RNAseq analysis. Proteolytic activity and pyochelin production were both increased, consistent with other studies which have shown that an increase in RpoN leads to a reduction in these phenotypes [37, 39]. The RM null mutant was less able to form biofilms under static conditions, both in microtiter plates and at the interface between culture medium and the air (pellicle). In *B. cenocepacia* K56-2, the RpoN sigma factor is required for bacterial motility and biofilm formation [50].

RM system acquisition occurred early in bacterial evolution [51]. The first investigations of RM systems demonstrated their important role in defence against foreign DNA by allowing self/non-self discrimination (reviewed in [34]). Conjugative transfer experiments to move megaplasmid pC3 between *B. cenocepacia* K56-2 and the RM system null mutant verified that RM systems indeed play an important role in protecting the *B. cenocepacia* H111 genome against incoming DNA, since in the absence of RM systems in the recipient such transfer increased 24-fold in efficiency.

The spontaneous loss of pC3 that occurred during the construction of the RM null mutant suggested to us that the deletion of RM components might have resulted in reduced genome stability. To quantitatively evaluate pC3 stability, the frequency of pC3 loss was determined using a modified version of an experiment previously described in [52]. This confirmed that RM systems and components play a central role in the maintenance of genome integrity in *B. cenocepacia* H111. The loss of pC3 occurred so frequently in the RM null mutant that separation of the modified null strain into pC3 deficient and positive strains was also observed through colony morphology. This reduction in stability explains why pC3 was spontaneously lost during the construction of the RM null mutant. In *E. coli* Dam methylase mutants, the timing of chromosome replication is disturbed, resulting in varying numbers of replicons in daughter cells [53]. Since there was little effect of the deletion of RM components and systems on H111 growth, however, we conclude that the effect on the stability of the essential replicons must be slight.

This study aimed to shed light on the involvement of DNA methyltransferases in the regulation of important cellular processes, as well as to unravel the impact of RM systems on bacterial phenotypes. Our work has confirmed the role of methylases and RM systems in genome protection and stability and has suggested involvement in phenotypes such as biofilm formation, siderophore production, motility, and prophage induction.

## Materials and Methods

### Bioinformatic analysis of RM components

A file containing the amino acid sequences of all RM components, both putative and experimentally proven, logged within the REBASE database was kindly provided by REBASE. An initial analysis was carried out on all *Burkholderia* and *Paraburkholderia* members within this file (due to the strain nomenclature used within REBASE, this meant that all *Burkholderia sensu lato* strain in the database were included). The translated sequences of the H111 RM genes (*gp10, I35_2397; gp51, I35_2438;* TIVRMc1, *I35_3250;* TIRE, I35_3252; TIM, I35_3254; TIIIRE, I35_1826; TIIIM, I35_1825; TIIM, *I35_2582;* TIVRMc2, *I35_ 1041*) were used as BlastP queries to find homologous components within the REBASE file using CLC Main Workbench v8. The percentage ID was calculated as the number of identical residues between the query and the match, as a percentage of the number of residues present in the H111 query sequence (excluding stop codon). This is shown in Fig S1.

A further analysis was carried out by downloading the genomes of a representative strain from each of the *Burkholderia sensu lato* species represented in REBASE. Where possible, a commonly studied type strain was chosen. Three *B. cenocepacia* strains were chosen to illustrate the diversity among some of the RM components within the species. For *Burkholderia fungorum*, no strain designation was listed in REBASE and strain ATCC BAA-463 was chosen. The same query sequences from the first analysis were used in a tBlastN search against these genome sequences. Percentage identity was calculated as described above. The phylogenetic tree was generated by concatenating the essential and highly conserved *gyrB* and *rpoD* genes from each species, aligning using CLC Main Workbench v8, trimming where less than 50 % of the sequences aligned with TrimaAl (Phylemon2), and then generating a phylogenetic tree using the neighbour joining method with CLC Main Workbench v8. The genome of *Ralstonia pickettiii* 12D was also included as an outgroup for the phylogenetic analysis.

### Bacterial strains, plasmids and media

All strains, plasmids and primers used in this study are listed in Tables S3 and S5 respectively. Unless otherwise stated, strains were grown aerobically in Luria–Bertani (Lennox) broth (Difco) at 37 °C. When required, media were supplemented with antibiotics at appropriate concentrations (in μg ml^-1^) as follows: chloramphenicol, 25 μg ml^-1^ (*E. coli*) and 50 μg ml^-1^ (Bcc); trimethoprim, 25 μg ml^-1^ (*E. coli*) and 50 μg ml^-1^ (Bcc); gentamicin, 20 μg ml^-1^ (*E. coli* and Bcc); and rifampicin, 50 μg ml^-1^ (Bcc). M9 medium containing uracil as the nitrogen source, as described previously [5], was used for differentiation between H111 and pC3 cured derivatives.

### Molecular techniques

Chromosomal DNA isolation was performed using the Wizard Genomic DNA Purification Kit from Promega, with minor modifications to the manufacturer’s protocol as follows. According to how much gDNA was required, a different amount of bacterial overnight culture was collected. After the cells were harvested by centrifugation, the pellet was resuspended in TNE buffer (10 mM Tris-HCL, 200 mM NaCl, 100 mM EDTA, pH 8) and incubated on ice for 20 to 30 min. The cell suspension was collected by centrifugation and the isolation protocol was carried out as per the manufacturer’s instructions. Plasmid preparation was routinely carried out using the Qiagen miniprep kit. DNA prepared by PCR amplification or restriction digestion was purified using the Qiagen PCR purification kit. Molecular methods were carried out as described by Sambrook et al. [54]. DNA fragments were amplified using either GoTaq DNA Polymerase (Promega) for diagnostic purposes or the proofreading Phusion High-Fidelity DNA Polymerase (NEB) to amplify fragments for use in cloning.

### Conjugal transfer of plasmids

Bacterial conjugations were used to introduce plasmids into Bcc strains, using a filter mating technique [55]. A helper strain (MC1061/pRK2013) was used to provide the *tra* genes. Conjugations were carried out on LB plates for approximately 16 h using saturated overnight cultures. *Pseudomonas* Isolation agar (PIA; Difco), supplemented with antibiotics as appropriate, was used for selection.

### Methylome Sequencing

Genomic DNA was extracted using the Wizard Kit (Promega), as stated above, and sequenced using Single Molecular, Real-Time (SMRT) sequencing on the PacBio RS II, by the Functional Genomics Center Zürich (FGCZ, University of Zurich). The raw data was analysed using the PacBio SMRT Portal. The sequenced reads were mapped to the reference sequence to allow detection of specific methylation patterns using the ‘Base Modification and Motif Analysis’ protocol.

### Data availability

The genomic data is available under NCBI BioProject number PRJNA609037. The raw reads of the sequenced genomic DNA are deposited in the SRA under the following accession numbers SAMN14218599 (*B. cenocepacia* H111 wild type), SRR11195332 (*B. cenocepacia* H111 null mutant), SRR11195331 (*B. ambifaria* AMMD), SRR11195330 (*B. multivorans* ATCC 17616), SRR11195329 (*B. lata* 383).

### Methylation visualisation

Prior to visualization, the abundance of modifications within each 10,000 bp length of DNA (window size) was calculated using ad-hoc Python scripts. The visualization of the detected modifications per window across the chromosomes was performed using the circlize package in R within the Rstudio interface version 1.1.463 [56]. We screened the genome for the presence of motifs in and around each replicons’ *oriC*, identified using the DoriC database, the oriFinder and DNAplotter ([57], [58], [59]).

### Verification of prophage region III loss in the RM system QM mutant

To investigate whether prophage region III was absent from the QM mutant, PCR was performed using primers designed to amplify genes within prophage region III encoding endolysin (gp12, I35_2399), holin (gp13, I35_4480) and the tail sheath protein (gp20, I35_2407). These genes could not be amplified from QM, but amplification was achieved from H111, and from each of the intermediate mutants leading up to QM. This suggested that phage region III had been lost in the construction of QM, and not in a previous step. We were able to amplify the genes flanking phage region III, suggesting that the phage had excised cleanly from the genome (data not shown). Clean loss of the prophage region III was later confirmed by sequence analysis of the null mutant (Fig S2, panel B).

### pC3 mobilization and curing

The mobilization of pC3 was enabled through insertion of an *oriT* via single crossover insertion of a suicide vector bearing an *oriT* (either pSHAFT2-gabD or pSHAFT2-araJ), allowing conjugative transfer, as previously described in [30]. To delete pC3 from several BCC strains, a straightforward replicon curing approach was performed using a constructed c3 mini-replicon called pMiniC3, bearing the single copy pC3 origin of replication, as described previously [30].

### pC3 stability assay

Assessment of pC3 stability was carried out as previously described in [52], with modifications to the protocol. Briefly, two specifically generated suicide vectors were used to construct strains to determine the pC3 stability. pEX18Gm-pMT-TpqueF carrying Tp resistance (*dhfrII*) under the regulation of a modified *lac* promotor was integrated into C1 via double homologous recombination. The gene encoding the repressor LacI was introduced into pC3 through a double crossover using pSHAFT2-nonconpJ23109-lacI-aacI. Strains were grown in IST media for 24 hour at 37 °C, and the cell count was determined by plating dilutions at intervals. Where pC3 was present in the cell, the expression of the Tp resistance gene was repressed by LacI. When pC3 was lost, Tp was expressed, resulting in colonies on IST plates containing Tp (25 μg ml^-1^). To test for spontaneous Tp-resistance, colonies were replica-plated on the pC3-selective medium M9ura [30] supplemented with Tp at 25 μg ml^-1^.

### Transfer efficiency test

To test for pC3 transfer efficiency between Bcc members, pC3 was mobilized by integration of pSHAFT2, which carries an *oriT*, allowing conjugative transfer, and a chloramphenicol resistance marker, in the donor strains (*B. cenocepacia* K56-2, *B. ambifaria* AMMD and *B. vietnamiensis* LMG 10929). The pC3 megaplasmid was cured from the recipient strains (*B. cenocepacia* H111 wild type, and RM system null mutant NullpC3^+^) as described by Agnoli and colleagues [30], and spontaneous rifampicin derivatives of the strains were selected by spreading 100 µl of the overnight culture on LB plates supplemented with 100 µg ml-1 rifampicin. Resistant colonies were restreaked on LB plates supplemented with rifampicin. Since the donor strains do not carry the *tra* genes required for formation of the sex pilus, a helper strain was used (MC1061/pRK2013), in a triparental mating. Dilution series were plated on PIA plates supplemented with Rif to calculate the total number of recipients and ex-conjugants). Depending on the strain, either 100 µl of an undiluted suspension or a dilution was plated on PIA plates supplemented with 200 µg ml^-1^ Cm and 50 µg ml^-1^ Rif to calculate the total number of ex-conjugants. Transfer efficiency was defined as total number of ex-conjugants/total number of recipients and ex-conjugants.

### Construction of conditional mutants

Conditional mutants were generated using the vector pSC200, which upon single crossover recombination with the genome separates a target gene from its native promoter, putting it under the control of the rhamnose-inducible PrhaB promoter [60]. Primers and restriction enzymes used have been detailed in Table S5. Conditional mutants were selected on PIA plates supplemented with 0.2 % rhamnose and trimethoprim (50 μg ml^-1^). To test for essentiality, conditional mutants were grown overnight in LB medium supplemented with 0.2 % rhamnose. Five µl from each sample of a dilution series was spotted on PIA media supplemented with either 0.5 % glucose or rhamnose for each strain. Plates were grown for 24 hours at 37 °C.

### Construction of targeted unmarked gene deletions

To construct markerless gene deletions, a protocol modified from that previously described by Flannagan was used [28]. Briefly, regions of homology of approximately 500 bp in size flanking the gene to be deleted were amplified using Phusion DNA Polymerase. Both fragments, as well as the vector pGPI-SceI, were digested with the chosen restriction enzymes. A tripartite ligation was performed, the plasmid transformed into electrocompetent *E. coli* SY327λpir and spread on selective LB plates. Positive clones were confirmed by colony PCR and sequence analysis using appropriate primers (see Table S5) and the plasmid introduced into *B. cenocepacia* H111 by triparental mating. Exconjugants were selected on PIA containing Tp and confirmed by PCR. A second homologous recombination was instigated by introducing vector pDAIGm-SceI into the recipient. Positive clones were selected on PIA plates containing gentamycin, verified by colony PCR and later colony purified by streaking on PIA plates without antibiotics. Primers used for the amplification and for the final deletion verification are stated in Table S5 and were designed using the H111 GenBank files (accession no. HG938370, HG938371 and HG938372).

The RM system null mutant was constructed by the sequential deletion of each RM region, resulting in a series of intermediate mutants, in addition to the final RM null mutant. The order of construction was as follows: 1) deletion of the 7054 bp Type I RM system (I35_3251 - I35_3254), encoded on C1, to give mutant TI; 2) deletion of the Type IV restriction endonuclease on C1 (I35_3250, 963 bp), to give strain DM; 3) deletion of the Type III RM system genes on C1 encoding one of the two core RM systems (I35_3273, I35_3274, 5051 bp), to give TM; 4) deletion of the Type IV restriction endonuclease on C2 (I35_ 5374, 729 bp), to give QM. Investigation of QM showed that prophage III had been lost from the genome, leaving only the Type II methylase gene (*I35_4914*) on C2. This was deleted to give NullpC3^-^. Upon discovery of the spontaneous loss of pC3 that occurred during the construction of NullpC3^-^, pC3 was moved back into the strain, as described in [7], to give NullpC3^+^. Finally, an unmarked RM null mutant strain was constructed by repeating the deletion of the Type II methylase gene (*I35_4914*) on C2 of QM, with care taken to select a pC3-containing clone. This strain was designated ‘Null’. The primers and restriction enzymes used for each deletion stage are indicated in Table S5.

### Preparation of samples for RNAseq analysis

Overnight cultures of the strains of interest were used to inoculate 50 ml LB broth with a starting OD_600_ of 0.01 and shaken at 220 rpm under aerobic conditions at 37 °C until an OD_600_ of 1 was reached. The culture was prepared and total RNA extraction carried out as detailed in [61]. To remove the remaining DNA, samples were treated with RQ1 RNAse-Free DNAse I (Promega) and purified using the RNAeasy MiniKit from QIAGEN, according to manufacturer’s guidelines. RNA quality was then checked with the RNA Nano Chip (Agilent 2100 Bioanalyzer; RNA Integrity Number >8) and 150 ng of total RNA were used for cDNA library construction. The Ovation Complete Prokaryotic RNA-Seq DR Multiplex System from NuGEN (NuGEN, San Carlos, CA, USA) was used to construct a strand-specific RNA-Seq library. This system uses Insert Dependent Adaptor Cleavage (InDAC) technology to remove ribosomal RNA. The cDNA library was analysed by capillary electrophoresis using a DNA chip from Agilent (Agilent High Sensitivity D1000 Screen Tape System). The prepared libraries were sequenced with the Illumina platform (single-end, HiSeq2500 instrument), by the Functional Genomics Center Zürich (FGCZ, University of Zurich). Between 6.2 and 9.5 million unique reads were obtained and mapped to the *B. cenocepacia* H111 genome using CLC Genomics Workbench v7.0 (QIAGEN CLC bio). The top 500 genes that showed the most significant changes in their expression (p-value ≤ 0.01 and absolute log_2_ (Fold change) ≥ 0.5) were taken for further analysis, and statistical analysis was performed using the *DESeq* R-package v1.26 [62]. The RNA-seq raw data files of wild type and mutant are accessible through the GEO Series accession number GSEXXXXXX.

### Swarming motility assay

Swarming motility was determined on nutrient broth plates containing 0.4 % agar, 0.5 % peptone and 0.3 % beef extract. Overnight cultures were normalized to an OD600 of 1, and 5 μl of the bacterial culture was spotted at the centre of the plate. After 24 hours of incubation at 30 °C, plates were documented photographically.

### Swimming motility assay

Swimming motility was measured on nutrient broth plates containing 0.3 % agar, 0.3 % peptone and 0.3 % beef extract. The plates were inoculated by touching the agar surface with a toothpick dipped into an OD_600_ 1 bacterial suspension and incubated for 24 hours at 30 °C.

### Colony morphology

Colony morphology was observed on NYG agar plates (1.5 % agar, 0.5 % peptone, 0.3 % yeast extract, and 2.0 % (w/v) glycerol). 5 µl of an overnight bacterial culture was spotted on the plates and incubated for 3 days at 37 °C, followed by a minimum of 2 days at RT.

### Biofilm formation assay

Biofilm formation was quantified in 96-well microtiter plates as described by ([63] and [64].The Biofilm Index (BI) was calculated as followed: BI= OD570 /OD550 * 100 [65].

### EPS production assay

EPS production was tested on YEM agar plates (0.05 % yeast extract, 0.4 % Mannitol, 1.5 % agar). Bacteria from an overnight culture were streaked and incubated for 48 hours at 37 °C.

### Pellicle formation assay

Pellicle formation was tested in NYG broth (0.5 % peptone, 0.3 % yeast extract and 2.0 % (w/v) glycerol). The media was inoculated 1:100 from a bacterial overnight culture and incubated at RT for a minimum of 5 days without shaking, in a capped tube to avoid evaporation.

### Antifungal activity assay

The antifungal activity assay was performed as previously stated in [30].

### Protease activity

Bacteria were assayed for proteolytic activity using the method of Safarik [66] with modifications to the protocol as described by Schmid and colleagues [67].

### Heat stress test

Bacterial overnight cultures were diluted to an OD_600_ of 1, and 500 µl was used to inoculate 50 ml LB broth (preheated to 42 °C). Cultures were incubated at 42 °C with 220 rpm shaking. The optical density was noted and the CFU µl^-1^ was monitored after 0, 3, 6 and 9 hours.

### Osmotic sensitivity assay

Resistance to osmotic stress was tested as previously stated in [52].

### Resistance to oxidative/chlorhexidine-induced stress

These assays were carried out as described by Kirby and colleagues [68], with modifications as described here. Whatman antibiotic assay discs (10 mm diameter) were used to test the resistance to two types of peroxide, inorganic (hydrogen peroxide; H_2_O_2_) and organic (*tert*-butyl-hydroperoxide), and the disinfectant agent chlorhexidine (which causes membrane disruption). Strains to be tested were grown overnight and the OD_600_ was adjusted to 1. LB plates were then inoculated in three planes using a cotton swab to give a bacterial lawn. 10 µl of 1 % *tert*-butyl-hydroperoxide, 2.5 % H_2_O_2_, or 20 % chlorhexidine were dropped onto discs (3 per strain/per replicate), placed on the inoculated plates and incubated overnight at 37 °C. Documentation was carried out either photographically or by measuring the diameter of the inhibition zone.

### *Galleria mellonella* pathogenicity assay

*Galleria mellonella* pathogenicity assays were carried out as described previously [7], using larvae purchased from BioSystems Technology, UK. Assays were performed in triplicate and 10 larvae were used per strain and control.

### Viability assay

PrestoBlue™ Cell Viability Reagent is a ready-to-use reagent, providing a quantitative measure of how metabolically active cells are. The assay was performed according to the manufacturer’s protocol. Samples of the cell suspension were taken at intervals over an incubation period of 24 hours and mixed with the reagent in a 1:10 ratio to a volume of 100 µl in a 96 well plate. To avoid unrepresentative results due to dye saturation, a dilution series was set up down the plate, with each well containing 10 µl reagent and 90 µl bacterial dilution. The mix was incubated for 1 hour at 37 °C and the fluorescence (Excitation: 530/25, Emission: 590/35) was measured in an MWGt Serius HT microplate reader from BioTek Instruments GmbH. To visualize viability over time, a non-saturated dilution (0.25) was chosen and plotted on a graph. In addition, optical density (OD_600_) of all replicates was measured, and the cell count was determined by plating dilutions at intervals on IST plates.

### Presto-blue cell viability test

For the cell viability test, the chloramphenicol markers present on pC3 in strains NullpC3^+^ and H111pC3^+^ were used to allow selection for pC3 maintenance. Strains were grown for 24 hours, with fluorescence measurements taken every three hours. Fluorescence after 24 hours was used to perform a two-tailed t-test.

### Use of fluorescence microscopy to examine cell morphology

Flasks with 20 ml LB broth were inoculated in duplicate with bacterial overnight cultures of NullpC3^+^ and H111pC3^+^ to a starting OD_600_ 0.01 and incubated at 37 °C with shaking. Samples were taken at time points 0, 4 and 24 hours after treatment and the plasma membrane was stained with FM 4-64 from Life Technologies (100 μg ml^-1^). Cells were observed with an epifluorescence Leica DM6000 B research microscope with a 100 x magnification.

### Phage visualization using transmission electron microscopy (TEM)

*Burkholderia cenocepacia* H111 and the RM null mutant (newNull) was cultured in 10 ml LB medium at 37 °C overnight. After centrifugation for 10 min at 5,000 rpm the supernatant was collected and filter-sterilized using a 0.22 µm pore size hydrophilic polyethersulfone filter (Merck Millipore, Germany). The cell-free supernatant was then ultracentrifuged at 150,000 x g for 1 hour. The pellet was resuspended in 50 µl PBS for visual phage detection. Phages or phage-like structures were absorbed on glow-discharged Formvar-coated 300-mesh copper grids and negatively stained with 1 % uranyl acetate for visualization using the Transmission electron microscopy (TEM).

### MV isolation and quantification

H111 wild type and the RM null mutant, Null, were grown overnight and used to inoculate flasks containing 20 ml LB broth to a starting OD_600_ of 0.02. Cultures were shaken for 24 hours at 37 °C. The isolation and quantification of the membrane vesicles was performed as described by Turnbull and colleagues [47]. Briefly, 10 ml of each bacterial suspension were spun for 10 min at 5,000 rpm (4472 rcf) at 4 °C, the supernatant collected and filter-sterilized using a 0.22 µm filter. After the supernatants were ultracentrifuged at 150,000 x g for 1 hour, each pellet containing membrane vesicles (MV) was resuspended in 100 µL PBS buffer. For quantification, MVs were stained with the membrane-binding dye FM1-43 (Life Technologies, USA) and the fluorescence intensity (510 nm excitation/626 nm emission) was measured using a MWGt Sirius HT microplate reader from BioTek Instruments GmbH.

### CAS assay

CAS plates were prepared to phenotypically characterize siderophore production, as described in [69]. 10 µl of each overnight culture was spotted on the plate and incubated for 48 hours at 37 °C. CAS plates were then visually inspected for a halo within and around the colonies, which indicated the production of iron chelating compounds, such as pyochelins.

### Pyochelin extraction and analysis

Strains were grown in 100 ml iron-free succinate (IFS) medium at 37 °C for 40 – 43 hours, until the OD_600_ was above 1. Cells were then collected by centrifugation for 20 min at 5000 rpm at 4 °C and the supernatant was sterile filtered, followed by acidification with 1 M HCl to a pH of 1.5 – 2 and extraction by adding 0.4 volumes of ethyl acetate. The upper ethyl acetate phase was then collected and vacuum dried using the Rotavapor RE (Büchi) until the amount was concentrated to about 5 ml total volume. This concentrate was then distributed into Eppendorf tubes and completely desiccated using the Eppendorf Concentrator 5301. The residue was then resuspended in 100 µl methanol and one µl of each sample analysed by thin layer chromatography (TLC) using silica gel 60 F254 (Merck Millipore, Germany) with chloroform-acetic acid-ethanol (90:5:10 [vol/vol]) as the developing solvent. The TLC plate was then quickly dipped into 100 mM FeCl3 to visualize the purified pyochelins, which were visible as brown areas on the TLC plate.

## Acknowledgements

We are grateful to Dr. Gabriella Pessi for her help with the RNA-seq analysis and for critical review of the manuscript. Thanks to Dr. Yilei Liu for her guidance in RNA library preparation. Many thanks to Dr. Anugraha Mathew and Dr. Yi-Chi Chen for their help with the chemical analysis and to Ratchara Kalawong for helping with the transmission electron microscopy. We would like to thank Dr. Carlotta Fabbri for excellent technical assistance. Many thanks also to Dr. Dana Macelis at REBASE, who at our request compiled a file containing the amino acid sequences of all RM components listed in the database, including their respective species and strains. This is available for ftp download and is updated on a monthly basis. PacBIO SMRT sequencing and Illumina sequencing for RNA-seq was carried out by the Functional Genomics Center Zürich (FGCZ, University of Zurich).

## Financial Disclosure

This work was supported by the Swiss National Fund (Project 31003A_122013, www.snf.ch) to LE. The funders had no role in study design, data collection and analysis, decision to publish, or preparation of the manuscript.

## Supporting information captions

**Figure S1. Homologues of the H111 RM components** within the *Burkholderia sensu lato.* A file containing the amino acid sequences of all RM components, both putative and experimentally proven, logged within the REBASE database was kindly provided by REBASE. The translated sequences of the H111 RM genes (*gp10, I35_2397; gp51, I35_2438;* TIVRMc1, *I35_3250;* TIRE, I35_3252; TIM, I35_3254; TIIIRE, I35_1826; TIIIM, I35_1825; TIIM, *I35_2582;* TIVRMc2, *I35_ 1041*) were used as BlastP queries to find homologous components within the REBASE file using CLC Main Workbench v8. The percentage ID was calculated as the number of identical residues between the query and the match, as a percentage of the number of residues present in the H111 query sequence (excluding stop codon).

**Figure S2. Distribution of modified bases in the RM null mutant.** The hatches surrounding each circle represent 10,000 bp windows. The number of modified bases per window is represented by the depth of colour. **A.** m4C modifications are shown in red, unassigned modified bases in grey. **B.** Distribution of ‘modified bases’ in the RM null mutant. Specific modifications could not be assigned, and probably represented DNA damage. The arrow indicates the area from which prophage region III was lost.

**Figure S3. Construction of conditional knockouts with a rhamnose inducible promotor confirmed the essentiality of *gp 51*.** OD_600_ adjusted cell suspensions of conditional *gp10* and *gp 51* mutants (CM10 and CM51, respectively) were spotted on medium supplemented with 0.5 % glucose or rhamnose. Growth was inspected after 24 hours at 37 °C.

**Figure S4. Growth of H111pC3^+^ and NullpC3^+^ swas similar.** Growth at 37 °C in LB medium was examined by determining OD_600_ over time.

**Figure S5. Phenotypic tests showing no significant impact by RM systems. A.** Plate assays for tests as labelled, from top to bottom: 1 % *tert*-butyl-hydroperoxide (organic peroxide, induces oxidative stress), 20 % chlorhexidine (induces stress to the membrane), EPS production on TEM plates, antifungal activity against *Rhizoctonia solanii.* **B.** Survival in medium containing 2M NaCl. **C.** Sensitivity to H_2_O_2_ (inorganic peroxide, induces oxidative stress). **D.** Survival at 42 °C. E. *Galleria mellonella* pathogenicity assay. Each image is representative of at least three biological replicates. Each graph shows the mean and SD for biological triplicates.

